# *In vivo* pharmacokinetics and tissue distribution profile of a Wnt/β-catenin pathway-targeting anticancer cassane diterpene isolated from *Caesalpinia pulcherrima*

**DOI:** 10.64898/2026.03.30.715187

**Authors:** Sandani De Vass Gunawardane, Oshadi V. Epitawala Arachchige, Shalini K. Wijerathne, P.A. Nimal Punyasiri, Arumugam Murugananthan, Sameera R. Samarakoon, Kanishka S. Senathilake

**Affiliations:** Institute of Biochemistry, Molecular Biology and Biotechnology, University of Colombo, No 90, Cumarathunga Munidasa Mawatha, Colombo 3, Sri Lanka; Department of Parasitology, Faculty of Medicine, University of Jaffna, Sri Lanka

## Abstract

A cassane diterpene, 6β-cinnamoyl-7α-hydroxyvouacapen-5α-ol (6βCHV), isolated from *Caesalpinia pulcherrima*, has emerged as a promising anticancer drug lead with reported Wnt/β-catenin pathway inhibitory activity and *in vivo* safety. The present study reports the *in vivo* pharmacokinetics and tissue distribution of 6βCHV in Wistar rats following a single oral dose of 200 mg/kg. A reproducible RP-HPLC-UV method was developed and validated for quantifying 6βCHV in rat plasma and tissues. Chromatographic separation was achieved using a gradient elution of methanol and water. The method was subsequently applied to investigate the pharmacokinetics and tissue distribution of 6βCHV. Plasma pharmacokinetic analysis revealed delayed and moderate absorption, with a T_max_ of 4 h and a C_max_ of 1314.12 ng/mL. Following absorption, 6βCHV is distributed widely across peripheral tissues, including the liver, heart, lungs, spleen, and kidneys, as well as pharmacological sanctuary sites such as the brain and testes. The highest concentrations were observed in the stomach, small intestine, and liver, with detectable levels persisting up to 24 h, reflecting extensive tissue partitioning and retention. Overall, these findings demonstrate that oral administration of 6βCHV is feasible. However, the delayed absorption suggests that further optimization of formulation or alternative administration routes may enhance systemic exposure. This study provides the first comprehensive pharmacokinetic and tissue distribution profile of 6βCHV, supporting its continued preclinical development as a potential anticancer therapeutic.

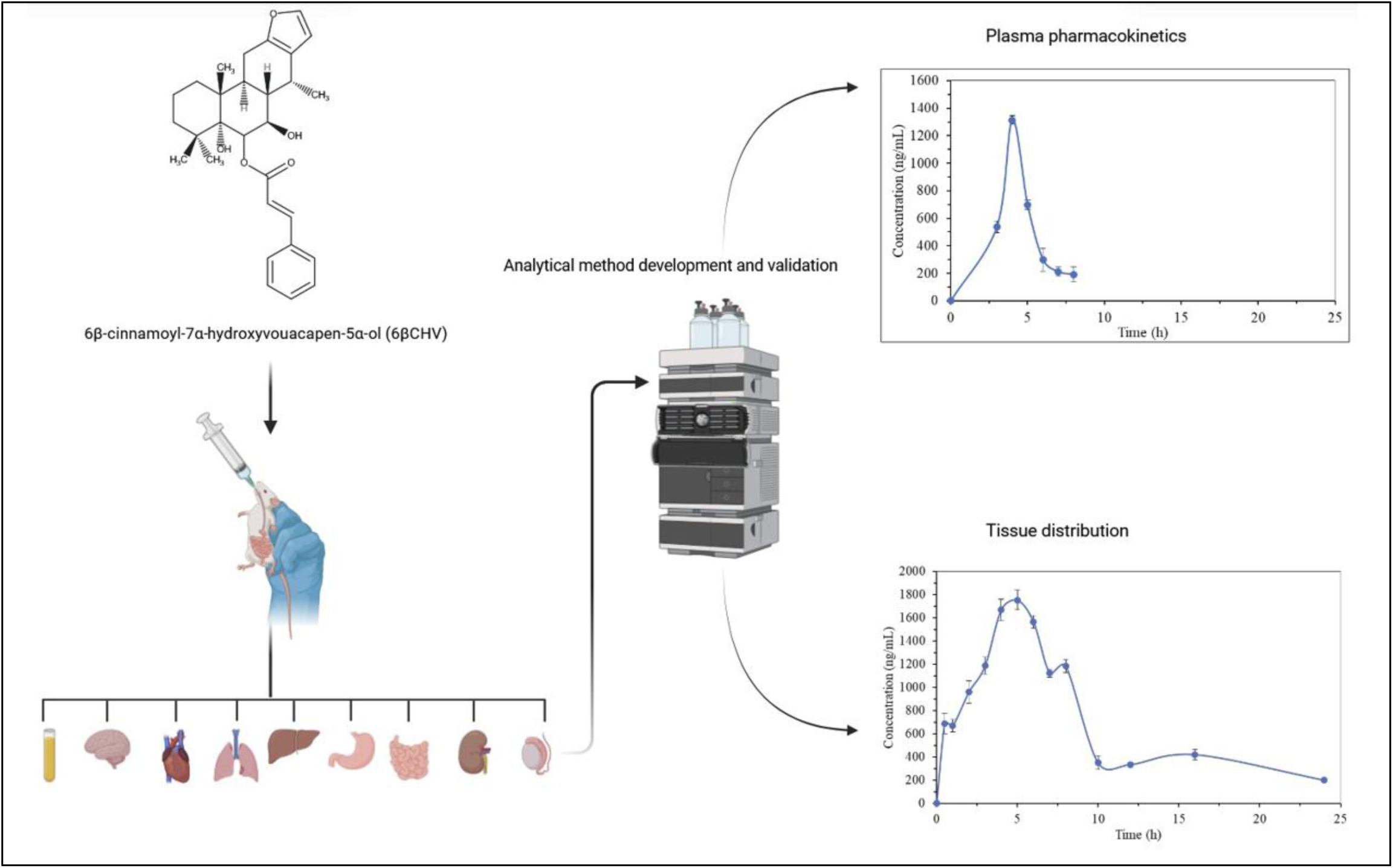

## 1. INTRODUCTION

Natural products remain a major source of structurally diverse and biologically active scaffolds for anticancer drug discovery. ^1^ Among them, diterpenes constitute a structurally diverse class of natural compounds composed of 20 carbon atoms derived from the condensation of four isoprene units. ^2^ The broad structural diversity of diterpenes underpins their diverse pharmacological activities, including anticancer, anti-inflammatory, antiviral, and immunomodulatory effects. Several diterpenes have been successfully translated into clinically used agents such as paclitaxel, oridonin, andrographolide, and ginkgolide, underscoring their value as lead compounds for cancer drug development. ^2–4^

Cassane diterpenoids, characterised by tricyclic cyclohexane rings fused to a furan ring, are prominent secondary metabolites of medicinal plants in the *Caesalpinia* genus. ^5^ These compounds have attracted increasing attention due to their potent cytotoxicity and favorable structure–activity relationships. Extracts and secondary metabolites from *Caesalpinia* species have demonstrated diverse anticancer activities, highlighting this genus as a promising reservoir of bioactive diterpenes. ^2,5,6^

The Wnt/β-catenin signaling pathway is frequently dysregulated in cancer stem cells (CSCs), ^7^ a subpopulation of tumor cells responsible for tumor initiation, metastasis, therapeutic resistance, and disease recurrence. ^8^ Despite its central role in cancer progression, therapeutic targeting of the Wnt/β-catenin signaling pathway remains challenging, and no small-molecule inhibitors have yet been approved for clinical use. ^9^ Recent computational screening of secondary metabolites from *Caesalpinia pulcherrima*, integrating molecular docking and molecular dynamics simulations, identified the cassane diterpene 6β-cinnamoyl-7α-hydroxyvouacapen-5α-ol (6βCHV) as a potent inhibitor of the β-catenin–Tcf/Lef interaction, a critical downstream event in Wnt/β-catenin signaling. Subsequent bioactivity-guided isolation and spectroscopic characterization confirmed 6βCHV as the major active compound in *C. pulcherrima* root, with promising antiproliferative activity against 16 cancer cell lines and CSC models (breast cancer stem cells and NTERA-2 cells). Notably, 6βCHV has been reported to exhibit higher antiproliferative activity against Wnt-dependent cancer types, including gastric adenocarcinoma, hepatocellular carcinoma, and ovarian carcinoma. ^10^

Furthermore, *in vivo* toxicity studies report that 6βCHV is well tolerated in Wistar rats following repeated dosing (15, 30, and 60 mg/kg for 28 days) and a single high oral dose at 300 mg/mL, with no toxicity observed, showing a safe *in vivo* toxicity profile. ^11^

Pharmacokinetic and tissue distribution studies are essential during the development of novel drug candidates. To reliably analyze complex biological samples, reversed-phase high-performance liquid chromatography with ultraviolet detection (RP-HPLC-UV) is widely employed in pharmacokinetic investigations due to its robustness, sensitivity, cost-effectiveness, and broad applicability to natural products in complex biological matrices. ^12^ To advance 6βCHV as a potential anticancer lead compound, a comprehensive understanding of its *in vivo* pharmacokinetic behavior is required, particularly with respect to absorption, distribution, metabolism, and elimination. However, despite its promising *in vitro* anticancer activity and established *in vivo* safety profile, neither a validated analytical method for detecting and quantifying the compound in biological matrices nor *in vivo* pharmacokinetic and tissue distribution profiles have been previously reported.

Therefore, in the present study, a novel and reproducible RP-HPLC-UV method employing propyl paraben as the internal standard (IS) was developed and validated for the first time for the quantification of 6βCHV in rat plasma and tissues. The validated method was subsequently applied to systematically characterize the plasma pharmacokinetics and tissue distribution of 6βCHV following oral administration in rats, representing the first comprehensive in vivo pharmacokinetic and tissue distribution evaluation of this compound. These data provide essential preclinical evidence to support the further development of 6βCHV as a promising anticancer lead.

## 2. RESULTS AND DISCUSSION

### 2.1. UV-Spectrum of 6βCHV

The UV spectrum of 6βCHV was recorded in the range of 200-300 nm, with a maximum absorbance (λ_max_) observed at 280 nm. Linearity was assessed based on absorbance values measured at 280 nm, and a correlation coefficient (R^2^) of 0.9994, suggesting good linearity. Figure 1 shows the UV spectrum and the corresponding linearity curve.

**Figure 1.**
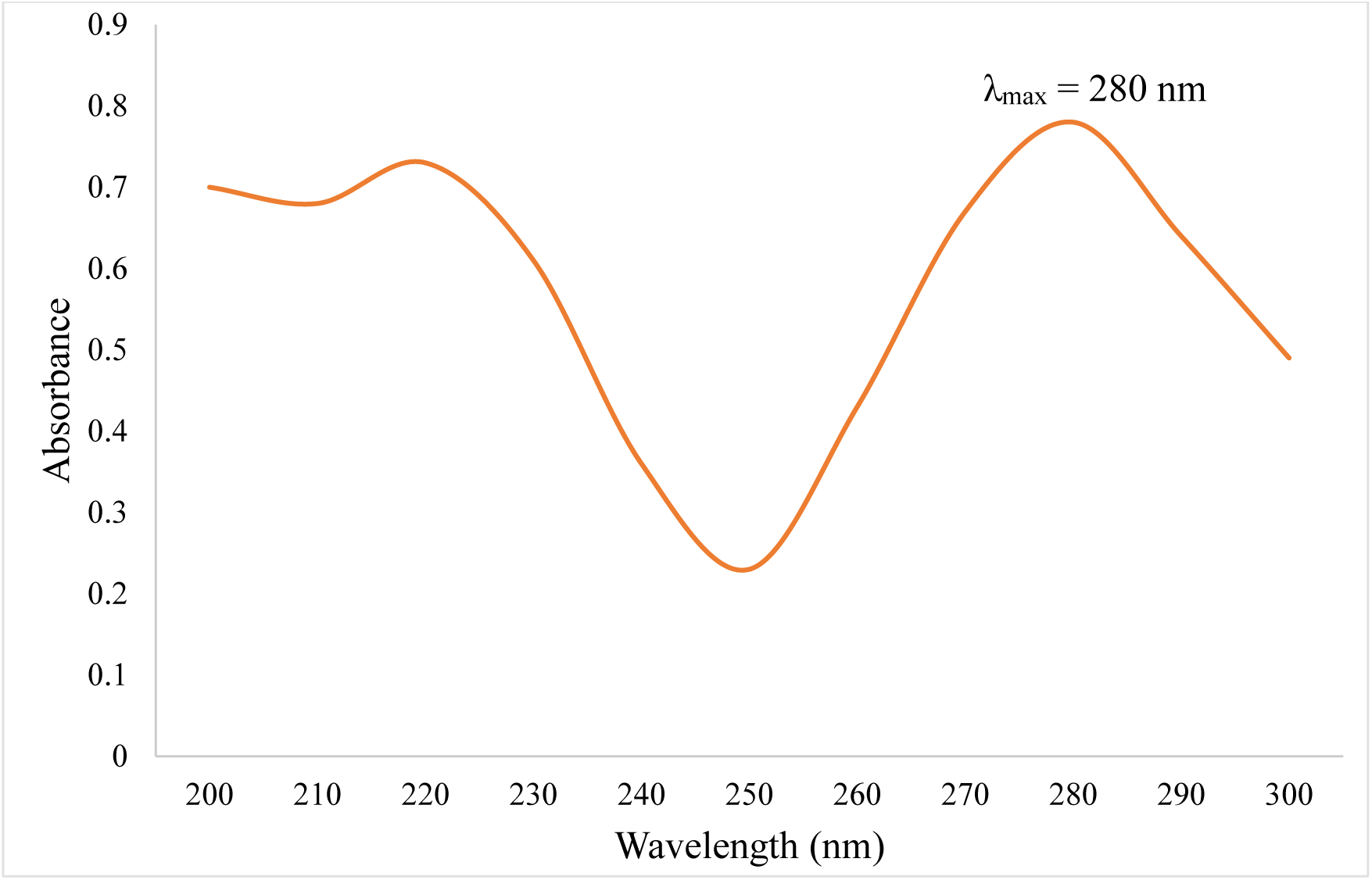
UV absorption spectrum of 6βCHV recorded in the range of 220 – 300 nm, showing a maximum absorbance (λ_max_) at 280 nm.

### 2.2. Optimization of Chromatographic Conditions

Chromatographic parameters were systematically optimized, including the column type, column temperature, UV detection wavelength, and mobile phase composition. Among several C18 columns evaluated, the Agilent Eclipse XDB-C18 column (4.6 mm × 150 mm, 5 µm) provided the best resolution and peak performance. The mobile phase composition was refined through multiple trials to achieve optimal resolution and symmetrical peak shapes for both 6βCHV and the IS. Various solvent systems, including methanol-water and acetonitrile-water mixtures at different ratios, were tested using both isocratic and gradient elution. A gradient elution program employing a methanol-water mixture with a variable flow rate over time yielded superior peak symmetry, enhanced separation, and reduced background noise. Propyl paraben (IS) exhibits strong UV absorbance within the 210-290 nm range. Following evaluation of several wavelengths, 275 nm was identified as the most suitable, providing optimal responses for both 6βCHV and propyl paraben. Under these optimized conditions, the retention times of 6βCHV and the IS were approximately 8 and 4 min, respectively.

### 2.3. Optimization of Sample Preparation

Efficient removal of plasma proteins and potential endogenous interferences is critical for the accurate quantification of analytes in biological matrices using HPLC. In the present study, various protein-precipitating agents, including methanol and acetonitrile with different ratios, were evaluated to optimize the sample preparation procedure. However, the extraction recoveries obtained with acetonitrile were unsatisfactory. Consequently, methanol-based protein precipitation (1:3, v/v) was selected due to its enhanced efficiency in protein removal and improved analyte recovery, and low background noise.

### 2.4. Internal Standard (IS)

After screening various compounds, propyl paraben was selected as the internal standard (IS) for method validation due to its stability, the absence of interference in biological matrices as an endogenous compound, and chromatographic conditions similar to 6βCHV (Figure 2). Moreover, propyl paraben demonstrated a strong UV absorbance in the range of 210–290 nm, with sufficient resolution from the analyte. Extraction recovery, matrix effect, and extraction efficiency of propyl paraben were also found to be highest when methanol was used as the protein precipitating agent.

**Figure 2.**
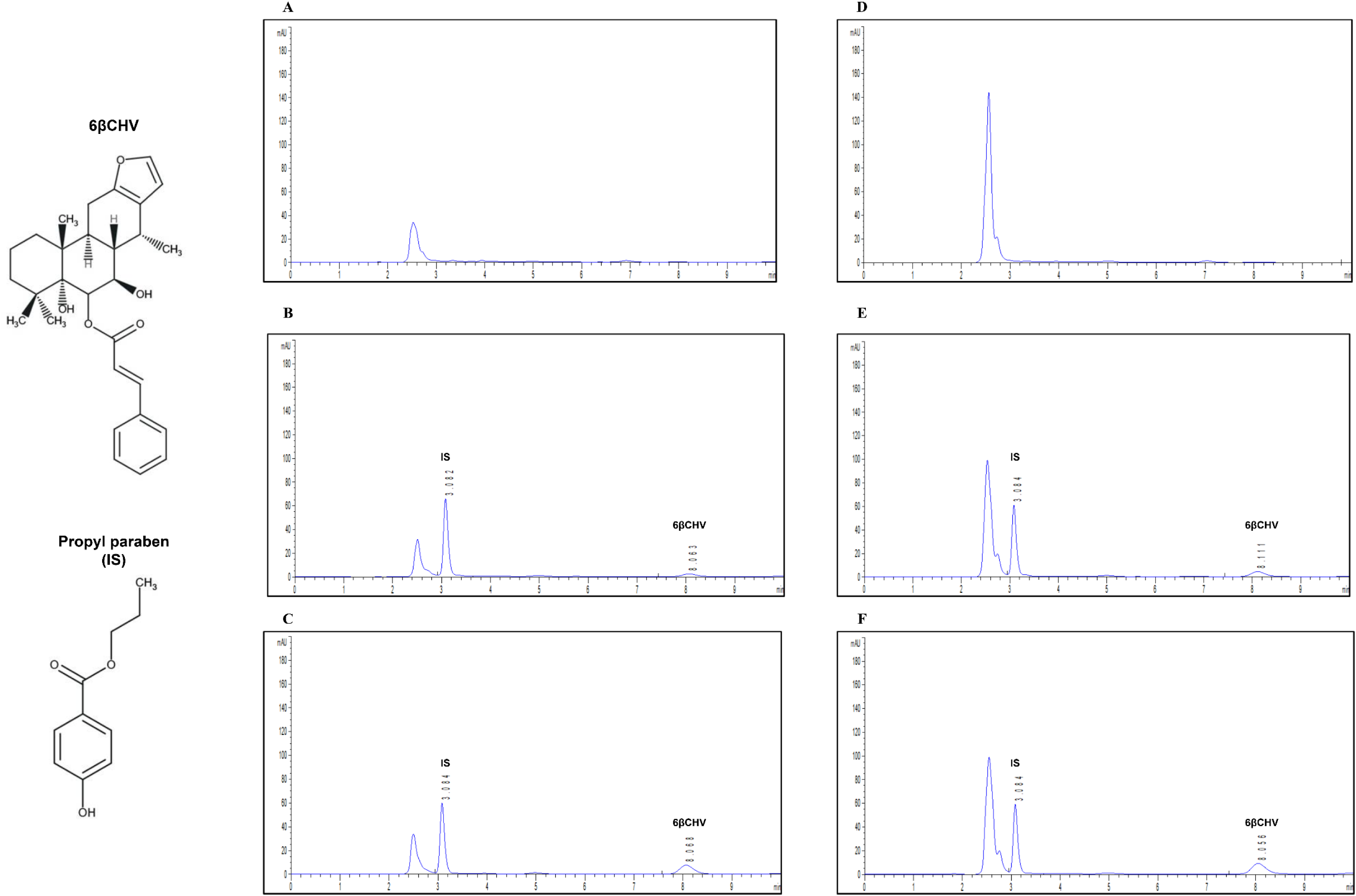
Representative chromatograms of rat plasma: (A) blank plasma sample, (B) plasma spiked with 6βCHV at LLOQ and IS, (C) plasma sample collected 4 h post-oral administration of 6βCHV (200 mg/kg), (D) blank liver sample, (E) liver spiked with 6βCHV at LLOQ and IS, and (F) liver sample collected 4 h post-oral administration of 6βCHV (200 mg/kg). IS: Internal standard; y axis: detector response (mAU); x axis: retention time (min). LLOQ: Lower limit of quantification; IS: Internal standard.

### 2.5. Method Validation

#### 2.5.1. Selectivity

The selectivity and specificity of the developed analytical method were evaluated by comparing the typical chromatograms obtained from blank matrices (matrix without analyte and IS), blank matrices spiked with analyte and IS, and samples from rats administered with 6βCHV. Interfering peaks were not observed at the retention times of 6βCHV or the IS in the blank matrices. Furthermore, chromatograms of blank matrices spiked with 6βCHV and IS confirmed the presence of clear and well-resolved peaks with no overlap from endogenous components. These findings indicate that the method was both selective and specific for the detection and quantification of 6βCHV in all tested biological matrices. Chromatograms of the blank matrix, spiked matrix, and samples from dosed rats are presented in the following figures, illustrating representative chromatograms from plasma (Figure 2A, B, C) and liver (Figure 2D, E, F).

#### 2.5.2. Linearity and Sensitivity

Calibration curves for 6βCHV were constructed by plotting the peak area ratios of 6βCHV to the internal standard against nominal concentrations for each biological matrix. All calibration curves demonstrated linearity over the validated concentration ranges, with correlation coefficients (R²) exceeding 0.9970 for plasma and all tissue matrices (Table 1), thereby surpassing the minimum acceptance criterion of 0.997 specified in the FDA guidelines for bioanalytical method validation. These results demonstrate a strong linear relationship between 6βCHV concentration and detector response across the concentration ranges relevant for pharmacokinetic and tissue distribution analysis. All back-calculated concentrations of calibration standards met the FDA acceptance criteria, confirming the method’s linearity over the tested concentration range (Table S1). Figure S1 present the calibration curves for 6βCHV in plasma and tissue matrices.

**Table 1.** The regression equations, correlation coefficients, linear range, and LLOQ for 6βCHV in plasma and tissue matrices (n = 3)

| Matrix | Calibration curve | Correlation coefficient ($R^2$) | Linear range (ng/mL) | LLOQ (ng/mL) |
| --- | --- | --- | --- | --- |
| Plasma | $y = 0.0001x - 0.0090$ | 0.9993 | 150 - 2550 | 150 |
| Liver | $y = 0.0001x - 0.0031$ | 0.9971 | 150 - 2550 | 150 |
| Small intestines | $y = 0.0001x + 0.0296$ | 0.9979 | 200 - 30000 | 200 |
| Stomach | $y = 0.0001x + 0.0370$ | 0.9977 | 200 - 30000 | 200 |
| Testes | $y = 0.0002x + 0.0342$ | 0.9985 | 150 - 5000 | 150 |
| Lungs | $y = 0.0002x + 0.0345$ | 0.9979 | 150 - 5000 | 150 |
| Kidneys | $y = 0.0002x + 0.0955$ | 0.9986 | 150 - 5000 | 150 |
| Brain | $y = 0.0002x + 0.0256$ | 0.9986 | 150 - 5000 | 150 |
| Heart | $y = 0.0002x + 0.0824$ | 0.9988 | 150 - 5000 | 150 |
| Spleen | $y = 0.0002x + 0.0736$ | 0.9987 | 150 - 5000 | 150 |

LLOQ: Lower limit of quantification

The sensitivity of the developed HPLC–UV method was evaluated by determining the LLOQ of 6βCHV in plasma and tissue homogenates. The LLOQ was defined as the lowest nonzero calibration level that could be quantified with acceptable accuracy (±20% RE) and precision (≤20% RSD). All LLOQ samples met the FDA bioanalytical method validation criteria across all matrices, confirming the suitability of the method for quantifying low concentrations of 6βCHV in biological samples.

#### 2.5.3. Accuracy and Precision

Intra- and inter-day accuracy and precision of the HPLC–UV method were evaluated at five QC levels for 6βCHV in plasma and tissue matrices. Intra-day accuracy and precision across all matrices ranged from −8.6% to 10.45% and from −0.67% to 11.95%, respectively, while inter-day accuracy and precision ranged from 0.24% to 11.26% and from 0.18 % to 19.05%, respectively (Table 2). All values were within the acceptable limits defined by FDA bioanalytical method validation guidelines, demonstrating that the method is accurate, precise, and reproducible for quantifying 6βCHV in all tested biological samples (Table S2).

**Table 2.** Intraday and interday accuracy and precision for 6βCHV in plasma and liver.

| Sample matrix | QC level | Nominal concentration (ng/mL) | Intraday (n = 30) |  | Interday (n = 25) |  |
| --- | --- | --- | --- | --- | --- | --- |
|  |  |  | Accuracy (RE%) | Precision (RSD%) | Accuracy (RE%) | Precision (RSD%) |
| Plasma | LLOQ | 150 | -5.90 | 3.18 | -6.77 | 1.41 |
|  | LQC | 450 | -0.67 | 2.70 | 0.85 | 2.08 |
|  | MQC | 850 | -1.95 | 0.51 | -1.78 | 0.49 |
|  | HQC | 2050 | 1.43 | 0.34 | 1.01 | 8.13 |
|  | ULOQ | 2550 | 0.55 | 0.14 | 0.34 | 0.18 |
| Liver | LLOQ | 150 | -8.33 | 2.98 | -5.35 | 2.90 |
|  | LQC | 450 | 0.15 | 6.74 | 2.25 | 3.90 |
|  | MQC | 850 | 1.91 | 4.33 | 1.13 | 1.13 |
|  | HQC | 2050 | 2.56 | 2.30 | -0.33 | 3.29 |
|  | ULOQ | 2550 | -1.12 | 1.41 | -1.25 | 1.50 |

LLOQ: Lower limit of quantification, LQC: Low quality control, MQC: Medium quality control, HQC: High quality control, ULOQ: Upper limit of quantification, RE: Relative error, RSD: Relative standard deviation.

#### 2.5.4. Extraction Recovery, Extraction Efficiency, and Matrix Effect

Extraction recovery, extraction efficiency, and matrix effects of 6βCHV and the internal standard (propyl paraben) were evaluated at three quality control (QC) levels (LQC, MQC, and HQC) in plasma and tissue matrices. Extraction recovery was determined by comparing peak areas of pre-extraction spiked samples with those of post-extraction spiked samples and ranged from 83.53% to 103.98% for 6βCHV across all matrices, indicating consistent and efficient extraction.

Extraction efficiency, assessed by comparing pre-extraction spiked samples with neat standard solutions, showed good reproducibility with values ranging from 80.81% to 97.90%.

Matrix effects were evaluated by comparing the peak responses of post-extraction spiked samples with those of neat standards. Matrix effect values for 6βCHV ranged from 83.00% to 102.07% across all matrices, indicating negligible ion suppression or enhancement. Detailed results for plasma and liver matrices are provided in Table 3, and Table S3 provides results for other tissue matrices.

**Table 3.**
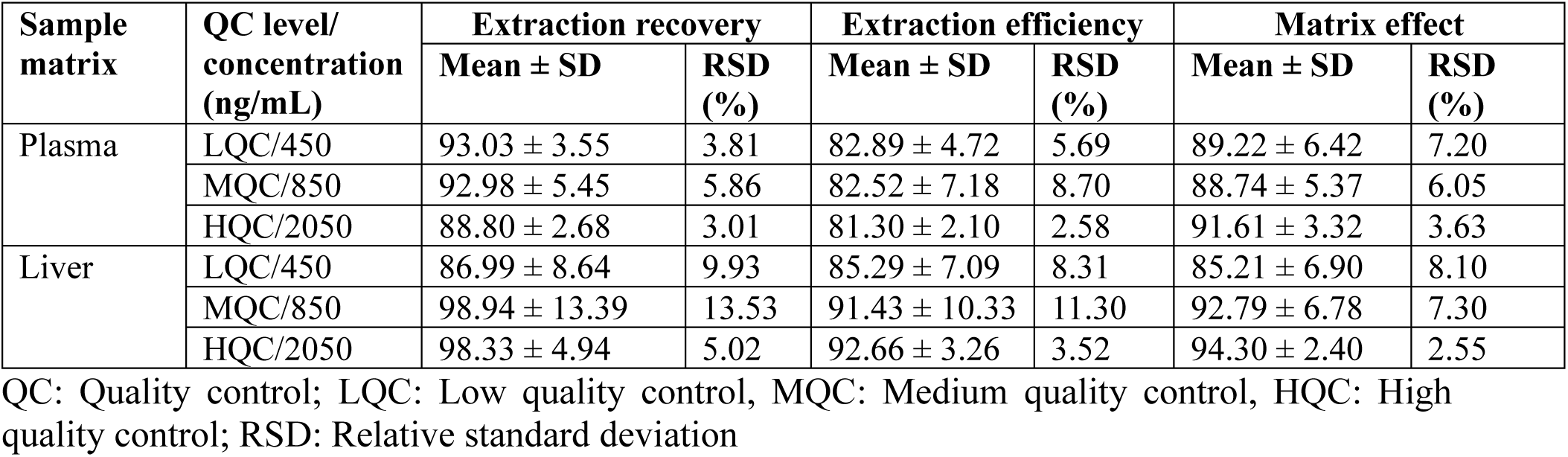
Extraction recovery, extraction efficiency, and matrix effect of 6βCHV in rat plasma and liver (n = 6)

| Sample matrix | QC level/<br>concentration<br>(ng/mL) | Extraction recovery |  | Extraction efficiency |  | Matrix effect |  |
| --- | --- | --- | --- | --- | --- | --- | --- |
| | | Mean ± SD | RSD (%) | Mean ± SD | RSD (%) | Mean ± SD | RSD (%) |
| Plasma | LQC/450 | 93.03 ± 3.55 | 3.81 | 82.89 ± 4.72 | 5.69 | 89.22 ± 6.42 | 7.20 |
| | MQC/850 | 92.98 ± 5.45 | 5.86 | 82.52 ± 7.18 | 8.70 | 88.74 ± 5.37 | 6.05 |
| | HQC/2050 | 88.80 ± 2.68 | 3.01 | 81.30 ± 2.10 | 2.58 | 91.61 ± 3.32 | 3.63 |
| Liver | LQC/450 | 86.99 ± 8.64 | 9.93 | 85.29 ± 7.09 | 8.31 | 85.21 ± 6.90 | 8.10 |
| | MQC/850 | 98.94 ± 13.39 | 13.53 | 91.43 ± 10.33 | 11.30 | 92.79 ± 6.78 | 7.30 |
| | HQC/2050 | 98.33 ± 4.94 | 5.02 | 92.66 ± 3.26 | 3.52 | 94.30 ± 2.40 | 2.55 |

QC: Quality control; LQC: Low quality control, MQC: Medium quality control, HQC: High quality control; RSD: Relative standard deviation

#### 2.5.5. Stability

6βCHV exhibited adequate stability in plasma and tissue homogenates under all tested storage and handling conditions (Table 4 and Table S4). At all three QC levels (LQC, MQC, and HQC), 6βCHV remained stable during short-term exposure at room temperature (6 h) and during autosampler storage at room temperature for 24 h, with measured concentrations remaining within acceptable limits. Long-term storage stability was also confirmed following 30 days at −20 °C for plasma and −80 °C for tissue homogenates. 6βCHV also maintained stability over three freeze–thaw cycles. These results indicate that sample handling, storage, and repeated freeze–thaw cycles do not compromise the integrity of 6βCHV, confirming the reproducibility of the method for routine pharmacokinetic and tissue distribution studies. In all cases, accuracy and precision values met FDA bioanalytical method validation criteria (±15%), indicating that sample handling and storage conditions did not compromise analyte integrity.

**Table 4.** Stability studies for 6βCHV in rat plasma and liver under different conditions.

| Sample matrix | QC level (ng/mL) | Short-term<sup>a</sup> |  | Long-term<sup>b</sup> |  | Freeze-thaw<sup>c</sup> |  | Autosampler<sup>d</sup> |  | Post-preparation storage<sup>e</sup> |  |
| --- | --- | --- | --- | --- | --- | --- | --- | --- | --- | --- | --- |
|  |  | RE% | RSD% | RE% | RSD% | RE% | RSD% | RE% | RSD% | RE% | RSD% |
| Plasma | LQC (450) | -7.54 | 2.39 | -8.12 | 2.12 | -7.19 | 3.01 | -11.57 | 5.85 | -8.66 | 3.01 |
|  | MQC (850) | -2.68 | 1.03 | -4.69 | 1.05 | 1.35 | -5.14 | -2.32 | 3.46 | -8.08 | 2.99 |
|  | HQC (2050) | -2.01 | 0.68 | -1.41 | 0.36 | -4.14 | 1.27 | -5.39 | 1.14 | -3.04 | 0.73 |
| Liver | LQC (450) | -9.06 | 2.13 | -7.92 | 2.40 | -10.25 | 3.21 | -5.26 | 5.36 | -10.82 | 3.96 |
|  | MQC (850) | -3.98 | 1.11 | -4.00 | 1.36 | 12.02 | 1.95 | -11.76 | 4.70 | -0.65 | 3.67 |
|  | HQC (2050) | -1.51 | 0.37 | -1.17 | 0.39 | -5.33 | 1.00 | -9.32 | 2.32 | -11.26 | 0.99 |

<sup>a</sup> Plasma and liver for 6h at room temperature. <sup>b</sup> Plasma and liver for 30 days at -20 °C and -80 °C, respectively. <sup>c</sup> Three freeze-thaw cycles from -80 °C to room temperature for liver and -20 °C to room temperature for plasma. <sup>d</sup> Plasma and liver samples were stored in the autosampler for 24 h before injection. <sup>e</sup> Injection-ready plasma and liver QCs at 4 °C for 48 h. LQC: Low quality control, MQC: Medium quality control, HQC: High quality control, RE: Relative error, RSD: Relative standard deviation.

#### 2.5.6. Carryover Effect and Dilution Integrity

Carryover was evaluated by injecting a blank plasma or tissue sample immediately following the ULOQ sample. No detectable peaks were observed at the retention times of 6βCHV (8.0 min) or the internal standard (4.0 min), confirming the absence of significant carryover. These results meet FDA acceptance criteria, which require that any response in the blank following the ULOQ does not exceed 20% of the LLOQ for the analyte and 5% for the internal standard.

Dilution integrity was assessed by analyzing plasma and tissue samples spiked with 6βCHV at concentrations above the ULOQ after a 2-fold dilution with blank matrix. Measured concentrations maintained accuracy within ±15% RE and precision ≤15% RSD, confirming that samples exceeding the validated range could be reliably quantified. Results for plasma and tissue matrices are summarized in Table S5, supporting the method’s suitability for handling samples above the ULOQ.

### 2.6. Plasma Pharmacokinetics

The newly developed and validated analytical method was applied to investigate the pharmacokinetics of 6βCHV in Wistar rats following a single oral dose of 200 mg/kg body weight. As shown in Figure 3, the maximum plasma drug concentration (C_max_) of 1314.12 ± 31.41 ng/mL was observed at 4 h (T_max_), indicating a delayed absorption profile. The relatively delayed T_max_ suggests a slow oral absorption of 6βCHV, which may be attributed to its high lipophilicity and poor aqueous solubility. Such characteristics are known to reduce the fraction of freely dissolved drug in the gastrointestinal lumen, thereby slowing the rate of absorption following oral administration. ^13^ This delayed absorption is comparable to other diterpenes reported in the literature, such as enmein, epinodosin, and isodocarpin. ^14^ The similarity in absorption kinetics suggests that delayed T_max_ reflects their lipophilicity and solubility-limited uptake in the gastrointestinal tract. Additionally, the use of corn oil as the dosing vehicle may have affected the absorption profile. Lipid-based vehicles such as corn oil are known to prolong residence time in the gastrointestinal tract and delay absorption, while enhancing the solubilization of lipophilic compounds, thereby shifting systemic uptake to later time points. ^15^

**Figure 3.**
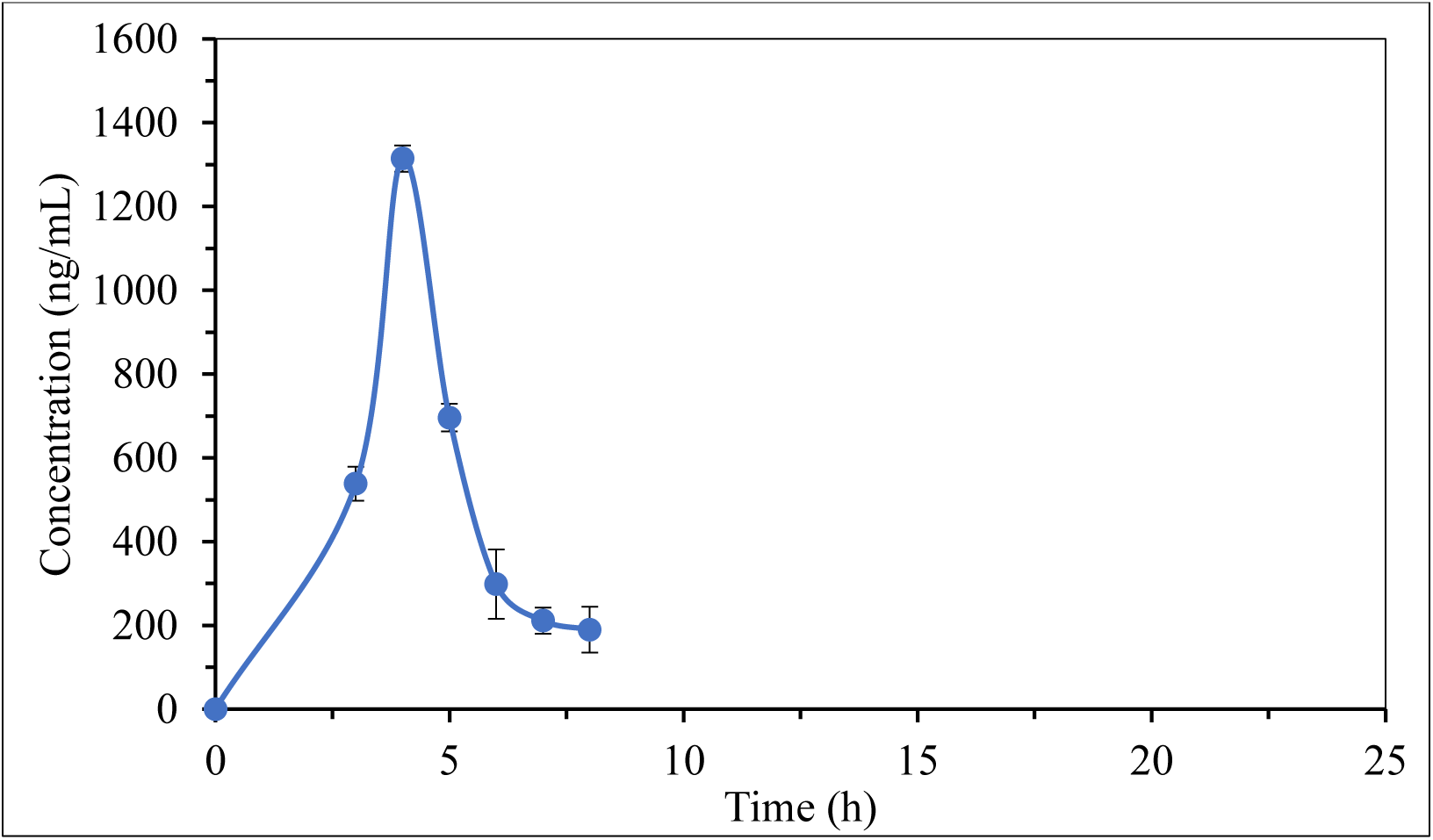
Plasma concentration-time profile of 6βCHV in Wistar rats following oral administration.

The systemic drug exposure, as measured by the area under the curve (AUC_0-*t*_), was 3690.89 ng/mL·h, suggesting a higher drug exposure despite the delayed rate of appearance in plasma. However, when normalized for dose, 6βCHV demonstrated lower systemic exposure per mg/kg than triptolide (a diterpenoid triepoxide) ^16^ or several other cassane diterpenes. ^2^ The T_1/2_ of 1.37 h indicates that 6βCHV is eliminated relatively quickly following oral administration. However, the mean residence time (MRT) in plasma was longer at 4.85 h, reflecting moderate systemic persistence despite the rapid decline in plasma concentrations. Compared to other diterpenes such as bonducellpin G, 7-O-acetyl bonducellpin C, and caesalmin E, which exhibit longer half-life and shorter MRT, ^2^ 6βCHV demonstrated faster elimination but sustained systemic exposure. This profile is consistent with triptolide and isodocarpin. ^14,16^

An apparent clearance (Cl/F) of 50 L/h/kg suggests a slow systemic elimination once absorbed. However, given the oral route, this clearance is normalized to bioavailability, and the modest plasma exposure despite the high dose of 6βCHV (200 mg/kg) suggests poor absorption and extensive first-pass metabolism rather than efficient systemic clearance. This is consistent with the high AUC values observed in the stomach and intestines, as discussed below. The low V_z_/F (0.10 mg/kg/(ng/mL)) suggests limited distribution beyond plasma when normalized to bioavailability. However, this systemic view is counterbalanced by the tissue distribution data of 6βCHV.

**Table 5.** Non-compartmental pharmacokinetic parameters of 6βCHV in rat plasma after single oral administration of 200 mg/kg of 6βCHV. Values represent calculated parameter values using the mean of n = 3 concentration values.

| Pharmacokinetic parameter | Value |
| --- | --- |
| C<sub>max</sub> (ng/mL) | 1314.12 |
| T<sub>max</sub> (h) | 4.00 |
| T<sub>1/2</sub> (h) | 1.37 |
| AUC<sub>0-t</sub> (ng/mL·h) | 3690.89 |
| AUC<sub>0-∞</sub> (ng/mL·h) | 4066.01 |
| MRT (h) | 4.85 |
| V<sub>z</sub>/F (L/kg) | 100 |
| Cl/F (L/h/kg) | 50 |

**Table 6.** Non-compartmental pharmacokinetic parameters of 6βCHV in rat organs following a single oral administration of 200 mg/kg. Values represent parameters calculated using the mean concentration obtained from three independent measurements (n = 3).

| Organ | Pharmacokinetic parameters |  |  |  |  |  |  |  |
| --- | --- | --- | --- | --- | --- | --- | --- | --- |
|  | C<sub>max</sub><br>(ng/mL) | T<sub>max</sub><br>(h) | T<sub>1/2</sub><br>(h) | AUC<sub>0-t</sub><br>(ng/mL·h) | AUC<sub>0-∞</sub><br>(ng/mL·h) | MRT<br>(h) | V<sub>z</sub>/F<br>(L/kg) | Cl/F<br>(L/h/kg) |
| Liver | 1752.41 | 5.0 | 6.1 | 15890.08 | 17654.15 | 10.73 | 100 | 110 |
| Stomach | 27549.02 | 0.5 | 4.3 | 139814.92 | 143958.31 | 6.19 | 80 | 1 |
| Small Intestines | 23377.15 | 3.0 | 4.6 | 126973.70 | 809920.36 | 6.21 | 10 | 2 |
| Testes | 2881.99 | 4.0 | 1.4 | 9171.96 | 10443.92 | 4.80 | 39 | 19 |
| Lungs | 1139.84 | 4.0 | 5.9 | 6519.57 | 8368.11 | 10.92 | 201 | 23 |
| Kidneys | 702.47 | 4.0 | 2.6 | 2351.20 | 3187.93 | 6.69 | 237 | 63 |
| Brain | 408.94 | 5.0 | 1.4 | 1482.62 | 1787.12 | 5.44 | 225 | 111 |
| Heart | 1807.90 | 4.0 | 3.6 | 5085.21 | 6176.47 | 6.72 | 166 | 32 |
| Spleen | 704.56 | 4.0 | 3.6 | 3980.72 | 4854.06 | 8.27 | 214 | 41 |

### 2.7. Tissue Distribution

Concentrations of 6βCHV were determined in the stomach, small intestine, liver, spleen, kidneys, heart, lungs, brain, and testes from Wistar rats at different post-dose time points. Figure 4 shows the concentration-time profiles of each organ over time, following oral administration of 6βCHV. The compound was quantifiable in all examined tissues, confirming a wide systemic distribution.

**Figure 4.**
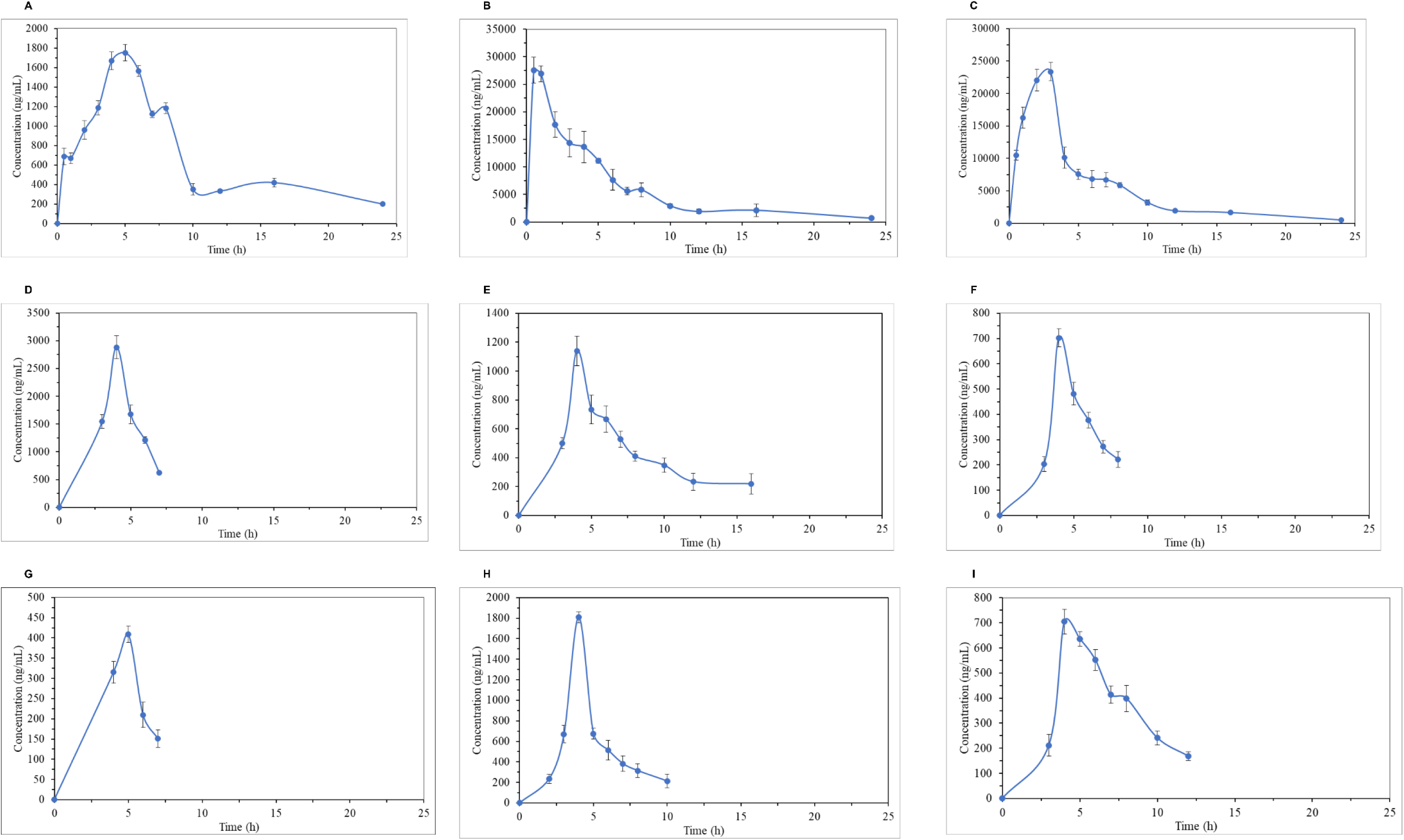
Concentration-time curve of 6βCHV in rat tissues following a single oral dose of compound (200 mg/kg). Data points represent the mean ± standard deviation (n = 3) for each time point. (A) Liver (B) stomach (C) small intestine (D) testes (E) lungs (F) kidneys (G) brain (H) heart (I) spleen.

As Figures 4B and 4C show, the gastrointestinal tract serves as the primary reservoir for 6βCHV, attributable to the oral administration route. Following gavage dosing, the compound is deposited directly into the stomach, resulting in an immediate, exceptionally high local concentration, with a C_max_ of 27549 ng/mL at 0.5 h. The C_max_ (23,377 ng/mL) and AUC (126,973.70 ng/mL·h) values observed in the small intestine, coupled with the gradual rise in plasma concentrations, support the interpretation that systemic absorption occurs predominantly through the small intestine. Moreover, the delayed T_max_ relative to the stomach (3 h vs. 0.5 h) reflects the time required for gastric emptying and subsequent intestinal uptake. Lipophilic drugs are primarily absorbed through the small intestine via passive diffusion, ^17^ and our findings for 6βCHV are consistent with previous studies reporting preferential intestinal uptake following oral administration. ^2,18^

As Figure 4A shows, 6βCHV reached the C_max_ of 1752.41 ng/mL at 5 h with a prolonged half-life of 6.1 h and MRT of 10.73 h, indicating sustained hepatic retention. The relatively high AUC and low apparent clearance suggest that once absorbed, 6βCHV undergoes extensive first-pass metabolism. This behavior parallels that of other lipophilic anticancer diterpenes, which also exhibit prolonged retention following oral administration. ^2^ Although hepatic accumulation of 6βCHV may enhance efficacy against primary liver cancers or hepatic metastases, it may also cause hepatotoxicity, as observed with anticancer agents such as capecitabine and lapatinib, both in combination therapies and as single agents. ^19–21^

The rank order of tissue drug exposure, as determined by AUC, was liver> testes> lungs> heart> spleen> kidneys> brain (excluding the stomach and small intestines), reflecting the combined influence of organ perfusion, tissue partitioning, and physiological barriers on distribution. Highly permeable compounds are known to distribute rapidly into well-perfused tissues and organs with fenestrated or discontinuous capillaries, such as the liver and spleen, which facilitates extensive tissue uptake and higher exposure. ^22^ The rank order of C_max_ followed a similar pattern, with the notable exception that the heart exhibited a higher C_max_ (1807.90 ng/mL) than the lungs (1139.84 ng/mL), despite the lungs (AUC_0-t_ = 6519.57 ng/mL·h) showing greater overall exposure than the heart (AUC_0-t_ = 5085.21 ng/mL·h). This divergence highlights the dynamic interplay between perfusion-driven uptake and tissue retention, where peak levels may not always correspond to exposure. ^22^ This interpretation is supported by the longer MRT in the lungs (10.92 h) compared with the heart (6.72 h) and the lower apparent clearance, indicating slower drug elimination from pulmonary tissue. Despite extremely high pulmonary perfusion and short diffusion distances, the distribution volume of the neutral and lipophilic compounds in lung tissue can be limited by relatively low lipid content, which may attenuate early peak concentrations. However, the generally enzyme-limited drug metabolism capacity of the lung favors prolonged retention rather than rapid clearance. ^23^ The relatively higher C_max_ observed in the heart reflects rapid perfusion-driven drug distribution. In contrast, the lower AUC, shorter MRT, and higher apparent clearance indicate limited tissue binding and rapid washout from myocardial tissue. Notably, high pulmonary AUC following oral administration has been reported for small molecules in previous studies. ^18,23^

In contrast, the kidneys exhibited both low C_max_ (702.47 ng/mL) and low AUC (2351.20 ng/mL·h), despite high renal perfusion. The relatively higher apparent clearance and moderate MRT indicate perfusion-driven transit and rapid elimination with minimal renal accumulation of 6βCHV. Similar renal exposure has been reported for several lipophilic compounds, reflecting a pivotal role of renal uptake and subsequent clearance. ^2,18,24^

Among pharmacological sanctuary sites, the testes exhibited substantial exposure (C_max_ = 2881.99 ng/mL; AUC _0–t_ = 9171.96 ng/mL·h), exceeding that of the heart, lungs, spleen, kidneys, and brain, with a T_max_ of 4 h. This indicates efficient systemic distribution despite the presence of the blood–testis barrier. This indicates efficient systemic distribution into the testes despite the presence of the blood-testis barrier, likely facilitated by the lipophilic nature of 6βCHV. Although the blood-testes barrier and drug transport network in the testes restrict the passage of many xenobiotics, ^24^ several other cassane diterpenes have been reported to distribute into testicular tissues. ^2^ Although cumulative exposure was higher, the shorter MRT and T_1/2_ suggest rapid uptake with limited long-term tissue retention. The lower apparent clearance (19 L/h/kg) relative to highly perfused organs further reflects barrier-regulated efflux and relatively low perfusion in testicular tissue.

The brain exhibited the lowest exposure among all tissues (C_max_ = 408.94 ng/mL; AUC_0-t_ = 1482.62 ng/mL·h), reflecting restricted penetration of 6βCHV across the blood-brain barrier. The delayed T_max_ suggests slow trans-barrier permeation, while the short T_1/2_ and the highest apparent clearance indicate rapid efflux mediated by active transport systems such as P-glycoprotein, which limit intracerebral drug accumulation. ^25^ Highly lipophilic anticancer agents, such as carmustine, readily penetrate the BBB. ^26^ However, efficient efflux and rapid turnover can still limit net brain exposure for many anticancer compounds. ^25^ Similar low cerebral accumulation has also been reported for several cassane diterpenes, ^2^ consistent with the distribution pattern observed with 6βCHV.

The broad tissue distribution of 6βCHV is pharmacologically advantageous for systemic cancer therapy. 6βCHV has been identified as a CSC–targeted lead that downregulates the Wnt/β-catenin signaling pathway, which is critical for CSC maintenance and self-renewal. ^10^ CSCs drive tumor progression, metastasis, therapeutic resistance, and disease recurrence and are known to occupy diverse tissue microenvironments beyond the primary tumor site. ^8^ Accordingly, the observed multi-organ distribution of 6βCHV supports its potential to access disseminated CSC populations. Importantly, measurable penetration into pharmacological sanctuary tissues, including the testes and brain, further strengthens its potential to target both primary and metastatic disease.

## 3. CONCLUSIONS

A reproducible RP-HPLC/UV method was developed and validated to quantify 6βCHV in rat plasma and tissues, enabling the first in vivo pharmacokinetic and tissue-distribution characterization of 6βCHV following oral administration. 6βCHV exhibited moderate systemic exposure with a delayed plasma T_max_, consistent with absorption-limited kinetics for a highly lipophilic compound, and demonstrated broad tissue distribution, including pharmacologically sanctuary sites. The moderate plasma exposure and the need for a relatively high dose indicate that further optimization of formulation and dosing strategies is warranted. Given the high lipophilicity of 6βCHV, advanced lipid-based delivery systems may enhance solubilization, intestinal absorption, and lymphatic transport, while parenteral routes may be explored to assess the impact of bypassing first-pass metabolism on systemic exposure and therapeutic efficacy.

## 4. MATERIALS AND METHODS

### 4.1. Chemicals and Reagents

The test compound 6βCHV used in this study was isolated and characterized as previously described by Senevirathne et al. ^10^ Purity (>98%) was verified by thin-layer chromatography, nuclear magnetic resonance (NMR) spectroscopy, and HPLC-UV analysis. The IS (propyl paraben) (purity ≥99%), HPLC-grade methanol, and pharmaceutical-grade corn oil (delivery vehicle for 6βCHV) were purchased from Sigma-Aldrich (Saint Louis, MO).

### 4.2. UV-Spectrum of 6βCHV

The UV absorption spectrum of 6βCHV was recorded using an Agilent Cary 60 UV-Vis spectrophotometer (Agilent Technologies, Santa Clara, CA, USA). A stock solution of 6βCHV (100 μg/mL) was prepared in methanol and appropriately diluted to obtain calibration samples (2-16 μg/mL). Spectral scans were performed using quartz cuvettes (1 cm) over the range 200-400 nm, with methanol as the blank. All measurements were carried out in triplicate, and the wavelength of maximum absorption (λ _max_) was determined. A calibration curve was constructed by plotting the mean absorbance at λ _max_ against concentration, and linearity was evaluated using the correlation coefficient (R^2^).

### 4.3. Instrumentation and Chromatographic Conditions

An Agilent 1260 Infinity series HPLC system (Agilent Technologies, Santa Clara, CA, USA) equipped with a quaternary pump delivery system (G1311C), robotic autosampler (G1329B), column thermostat (G1316A), and multi-wavelength UV detector (G1315D). The chromatographic separation was implemented by an Agilent Eclipse XDB C18 column (5 µm particle size, 100 Å pore size, L × I.D. 15 cm × 4.6 mm, Agilent Technologies, Palo Alto, CA, USA). To enhance the separation and resolution of 6βCHV and IS from potential interfering substances in the plasma and tissue matrices, a gradient elution method with variable flow rate was employed. The mobile phase was comprised of a mixture of A (methanol) and B (water). The program of gradient elution was as follows: 0.00 min, 70% A and 30% B at 0.9 mL/min; 3.00 min, 80% A and 10% B at 0.4 mL/min; 5.00 min, 90% A and 10% B at 0.4 mL/min; 7.00 min, 100% A at 0.4 mL/min, 15 min, 70% A and 30% B at 0.9 mL/min; 20.00 min, 70% A and 30% B at 0.9 mL/min. The total chromatographic separation time was 20 min. The results were analyzed using OpenLab CDS EZChrom software. The column and autosampler temperatures were maintained at 25 °C and room temperature, respectively. The UV detection was carried out at 275 nm, and the injection volume was 20 µL.

### 4.4. Animals

All animal care and experimental procedures were approved by the Ethics Review Committee of the Institute of Biologists, Sri Lanka (IOBSL) (ERC IOBSL 384/12/2024), prior to the start of the experiments and were conducted in accordance with the national and institutional guidelines.

Forty-eight healthy male Wistar rats (*Rattus norvegicus*) (200-220 g) were purchased from the Medical Research Institute (MRI) of Sri Lanka. All animals were housed in standard polycarbonate cages bedded with sterile wood shavings to a depth of 2 cm. Animals were acclimatized for seven days under controlled environmental conditions (temperature: 24 °C ± 0.5 oC, humidity: 47.5 ± 2.5%, and a 12/12-hour light/dark cycle). The diet consisted of standard rat chow and water *ad libitum*. Following acclimatization, forty-two animals were randomly divided into fourteen groups (n = 3 per group) for pharmacokinetic and tissue distribution studies, and six rats were allocated for method validation. Before executing the pharmacokinetic and tissue distribution studies, the animals were fasted for 12 h.

### 4.5. Plasma Sample Extraction and Preparation

Plasma samples were extracted by an optimized protein precipitation method. A total of 95 μL of plasma was transferred to a 1.5 mL Eppendorf tube, 5 μL of the IS working solution was added, and vortexed for 1 min. Proteins in the matrix were precipitated by the addition of 300 μL of HPLC-grade methanol and vortexed for 1 min. Samples were then centrifuged at 14,000 rpm for 10 min at 4 °C. The supernatant was transferred to a new Eppendorf tube and evaporated to dryness under a 37 °C mild stream of nitrogen. The dried residue was then reconstituted by adding 100 μL of the initial mobile phase, a mixture of methanol and water (70:30, v/v), and vortexed for 1 min. The reconstituted samples were again centrifuged at 14,000 rpm for 10 min at 4 °C to remove any residual particulates. The supernatant was transferred to a new 1.5 mL Eppendorf tube, filtered through a nylon syringe filter (0.22 μm pore size, 13 mm diameter) into a new HPLC autosampler vial, and injected onto the HPLC system.

### 4.6. Tissue Sample Extraction and Preparation

Tissue samples were extracted using the same protein precipitation method used for plasma. Frozen rat organs were thawed, weighed, and homogenized using a mechanical homogenizer in ice-cold physiological saline (1:3 w/v ratio). ^14^ An aliquot of 190 μL of tissue homogenate was transferred to a 1.5 mL Eppendorf tube, followed by the addition of the IS (10 μL from 100 μg/mL), and vortexed for 1 min. The samples were then deproteinized with 600 μL of methanol, subjected to the subsequent sample processing step as plasma samples, and analyzed.

### 4.7. Analytical Method Validation

A complete validation of the bioanalytical method was carried out following the US Food and Drug Administration (FDA) guidelines for bioanalytical method validation and study sample analysis (Version 2022), in terms of selectivity, linearity, sensitivity, accuracy, precision, extraction recovery, matrix effect, carryover, and dilution integrity. ^27^

#### 4.7.1. Preparation of Stock Solutions, Calibration Standards and Quality Controls

The stock solutions of 6βCHV and IS were prepared at a concentration of 1.0 mg/mL in HPLC-grade methanol and stored in the dark at -20 °C The IS stock solution was further diluted with methanol to obtain the working solution at a concentration of 100 μg/mL. The stock solution of 6βCHV was appropriately diluted to prepare working solutions in the range of 1.5-50 μg/mL. Calibration samples were prepared by spiking the respective matrix with a series of known concentrations of 6βCHV. The calibration range was defined by the lower limit of quantification (LLOQ) and the upper limit of quantification (ULOQ).

Calibration samples for plasma were prepared by spiking 85 μL of blank plasma with 10 μL of the respective 6βCHV working solutions to achieve concentrations of 150, 250, 450, 650, 850, 1050, 1550, 2050, and 2550 ng/mL. For tissue matrices, calibration series were prepared by spiking 170 μL of blank tissue homogenate with 20 μL of the respective working solution to obtain the following concentration: 150, 250, 450, 650, 850, 1050, 1550, 2050, and 2550 ng/mL for liver; 200, 500, 1000, 2000, 5000, 10000, 15000, 20000, 25000, and 30000 ng/mL for small intestines and stomach; 150, 250, 450, 650, 850, 1050, 3050, 4050, and 5000 ng/mL for testes, lungs, kidneys, brain, heart, and spleen. The IS working solution was added to each calibration sample (5 µL for plasma and 10 µL for tissues) to a final concentration of 5000 ng/mL, followed by extraction, as described above. Calibration samples were daily prepared during the ongoing analysis.

Quality control (QC) samples were prepared in the same manner to obtain five QC levels, including LLOQ, low quality control (LQC), medium quality control (MQC), high quality control (HQC), and ULOQ. QC samples at aforementioned levels were prepared at the following concentrations: 150, 450, 850, 2050, and 2550 ng/mL for plasma and liver; 200, 1000, 10000, 20000, and 30000 ng/mL for stomach and small intestines; 150, 450, 1050, 3050, and 5000 ng/mL for testes, kidneys, brain, lungs, heart, and spleen.

#### 4.7.2. Selectivity and Specificity

Selectivity was evaluated by comparing chromatograms of blank matrix obtained from six rats, blank matrix spiked with 6βCHV at the LLOQ and IS, and the respective matrix sampled at 4 h after oral administration of 6βCHV. Selectivity of the analytical method was considered acceptable if no significant peaks were observed at the retention times of 6βCHV and the IS. Responses in the blank matrix attributable to interfering components should not be more than 20% of the 6βCHV response at the LLOQ, and not more than 5% of the IS response in the LLOQ sample for each matrix.

Specificity was assessed by evaluating chromatograms of the above samples to confirm that the analytical signal at the retention time was solely attributable to 6βCHV and not confounded by other interferences from endogenous matrix components. The response at the retention time of 6βCHV in the blank matrices should not exceed 20% of the 6βCHV response at the LLOQ and not more than 5% of the IS response in the LLOQ sample.

#### 4.7.3. Linearity and Sensitivity

The calibration curves were constructed by plotting the peak area ratios of 6βCHV to the IS (*y*-axis) against the nominal concentration of 6βCHV (*x*-axis). Linearity was assessed within the calibration range by fitting the data to a linear regression model and evaluating the correlation coefficient (R^2^ ≥0.99). The sensitivity was determined by analyzing six independent replicates of the LLOQ of each calibration curve. The LLOQ was required to demonstrate an accuracy within ±20% of the nominal value and precision not exceeding ±15%.

#### 4.7.4. Accuracy and Precision

The intraday accuracy and precision were determined by comparing five replicates of QC samples at the LLOQ, low (LQC), medium (MQC), high (HQC), and ULOQ concentration levels in each matrix with their nominal concentrations in a single batch. The interday accuracy and precision were measured by comparing five QC concentrations (five replicates per QC level) from three independent batches run on three consecutive days. The accuracy and precision of the method were expressed as percent relative error (RE%) and relative standard deviation (RSD%), respectively. The LLOQ was defined as the lowest concentration that could be determined with both RE and RSD within ±20%.

#### 4.7.5. Extraction Recovery, Extraction Efficiency and Matrix Effect

Extraction recovery, extraction efficiency, and matrix effect were determined at LQC, MQC, and HQC across all matrices by comparing the peak areas of the analyte between three types of samples: blank biological matrix spiked with QCs before sample processing (BE), blank biological matrix spiked with QCs after sample preparation (AE), and matrix-free solvent samples spiked with QCs (C). Quality control samples were prepared using matrices from six untreated rats. The following equations were used for calculating the extraction efficiency, the extraction recovery, and matrix effect. ^28^

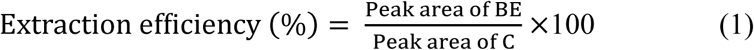

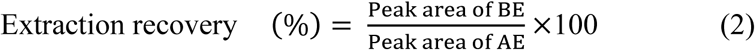

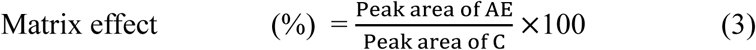

**Equation 1**: Equations to calculate extraction efficiency (1), Extraction recovery (2), and Matrix effect (3). BE: Pre-extraction spiked; AE: Post-extraction spiked; C: Neat solution in methanol

#### 4.7.6. Stability

The stability of 6βCHV in plasma and tissues was measured in six replicates at three QC levels (LQC, MQC, and HQC) under the following conditions. The short-term stability of 6βCHV was assessed by storing the QC sample on the laboratory benchtop at room temperature for 4 h. Long-term stability of 6βCHV in plasma and tissue matrices was evaluated by storing the QCs at -20 °C and -80 °C, respectively, for 30 days before analysis. Freeze-thaw stability of 6βCHV was assessed by subjecting QCs to three freeze-thaw cycles: plasma QCs were cycled between -20 °C and 25 °C, and tissue QCs between -80 °C and 25 °C. Ready-to-inject QCs were stored in the autosampler for 24 h to assess the post-preparative stability of 6βCHV in plasma and tissues.

#### 4.7.7. Carryover Effect and Dilution Integrity

The carryover effect was assessed by injecting each blank matrix immediately after the highest calibration sample (ULOQ) for that matrix. The study samples may contain 6βCHV concentrations outside the calibration range. Therefore, the effect of dilution was evaluated by preparing dilution QCs, which contain 6βCHV at a concentration higher than the ULOQ, and then diluting with the respective blank matrix to bring the concentration within the validated calibration range. Diluted QCs were prepared in five replicates at a 2-fold dilution level: 3000 and 5100 ng/mL for plasma and liver; 40000 and 60000 ng/mL for stomach and small intestines; 7000 and 10000 ng/mL for testes, lungs, kidneys, brain, heart, and spleen.

### 4.8. Pharmacokinetic and Tissue Distribution Studies

The validated analytical method was subsequently applied to investigate the pharmacokinetic and tissue distribution profile of 6βCHV following oral administration of a single dose of 200 mg/kg at a 5 mL/kg dosing volume, selected based on previous *in vivo* toxicity studies. ^11^ The compound was prepared for oral administration by dissolving the compound in pharmaceutical-grade corn oil, followed by vortexing for 5 min and sonication at 37 °C for 10 min to ensure complete dispersion and homogeneity before administration. Following overnight fasting, rats (n = 3 per time point) were orally administered the prepared 6βCHV suspension, and fasting was continued for an additional 4 h. Subsequently, animals were deeply anesthetized by diethyl ether inhalation and euthanized at pre-dose (0 min) and 0.5, 1, 2, 3, 4, 5, 6, 7, 8, 10, 12, 16, and 24 h post-dose by terminal cardiac puncture exsanguination. Blood was immediately collected (∼3.0 mL) and transferred into EDTA tubes on ice. After centrifuging at 5000 rpm at 4 °C for 10 min, plasma was recovered and stored at -20 °C until analysis. Following blood collection, the organs, including the brain, heart, lungs, liver, kidneys, spleen, stomach, small intestines, and testes, were immediately collected. Organs were rinsed with ice-cold physiological saline to remove blood and contents, gently blot-dried, weighed, and snap-frozen in liquid nitrogen to immediately stop enzymatic activity. The organs were then stored at -80 °C until processing and analysis.

#### 4.8.1. Pharmacokinetic and Tissue Distribution Data Analysis

All experimental data are presented as the mean ± standard deviation (SD) of measurements from three rats at each time point. Data analysis was performed by non-compartmental analysis using PKSolver 2.0 (Microsoft Excel add-in). Pharmacokinetic parameters determined included the area under the concentration-time curve from time zero to last measurable concentration (AUC_0-t_), the area under the concentration-time curve from zero to time infinity (AUC_0-∞_), mean residence time (MRT), half-life (t_1/2_), time of maximum observed concentration (T_max_), peak concentration (C_max_), apparent volume of distribution (V_z_/F), apparent body clearance (CL/F) for both plasma and tissue matrices.

## Funding Sources

### Notes

The authors declare no competing interests.

