## Supporting Information - Figures for "*In vivo* pharmacokinetics and tissue distribution profile of a Wnt/β-catenin pathway-targeting anticancer cassane diterpene isolated from *Caesalpinia pulcherrima*"

**Figure S1. Calibration curves of 6BCHV in rat (A) plasma, (B) liver, (C) small intestines, (D) stomach, (E) testes, (F) lungs, (G) kidneys, (H) brain, (I) heart, and (J) spleen.**

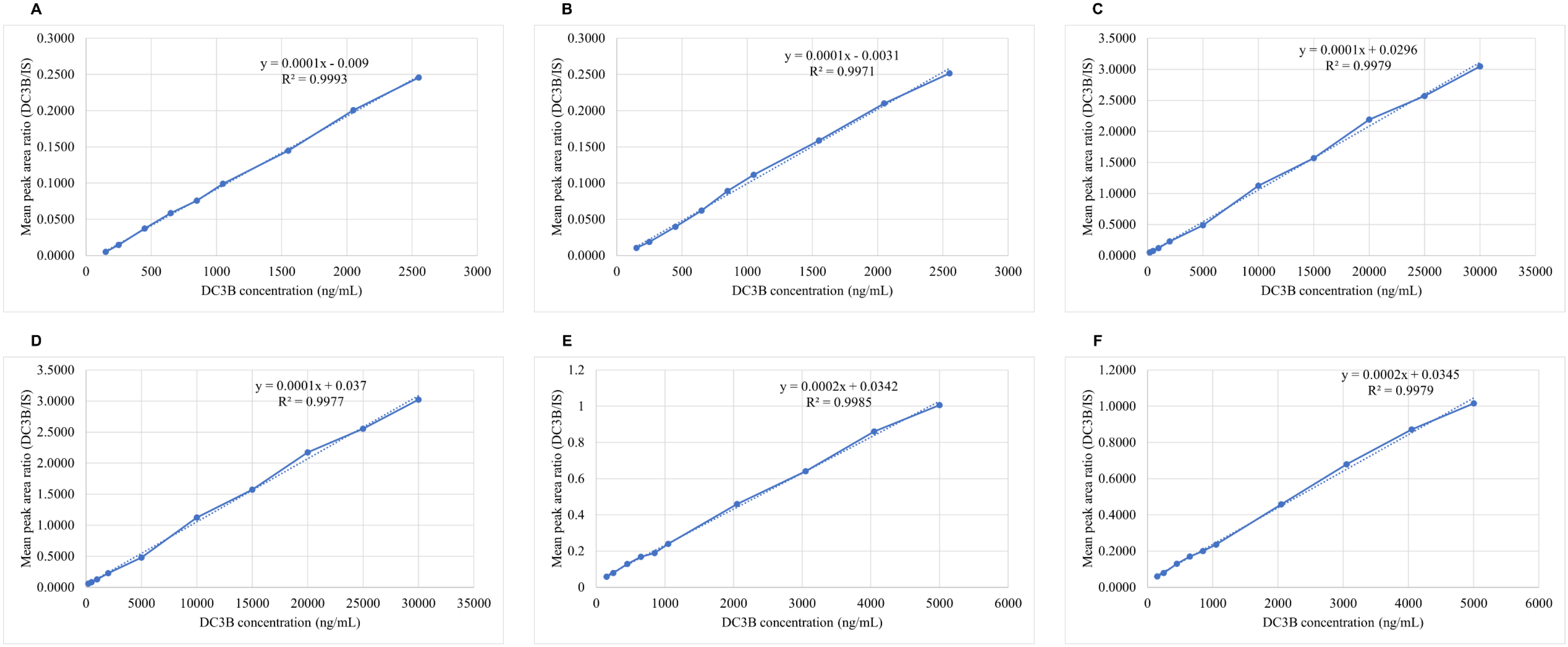

**G**

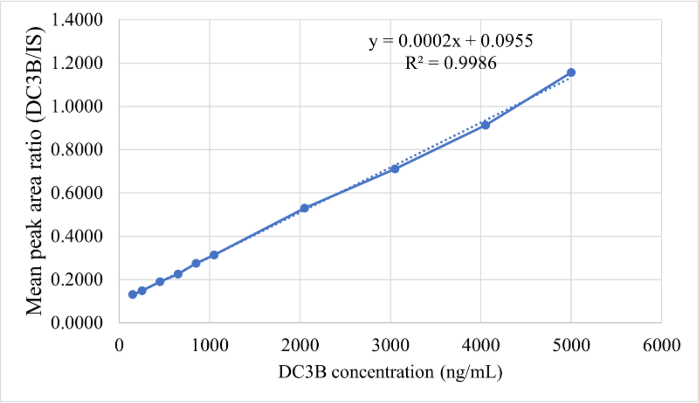

y = 0.0002x + 0.0955
R² = 0.9986
Mean peak area ratio (DC3B/IS)
DC3B concentration (ng/mL)


**H**

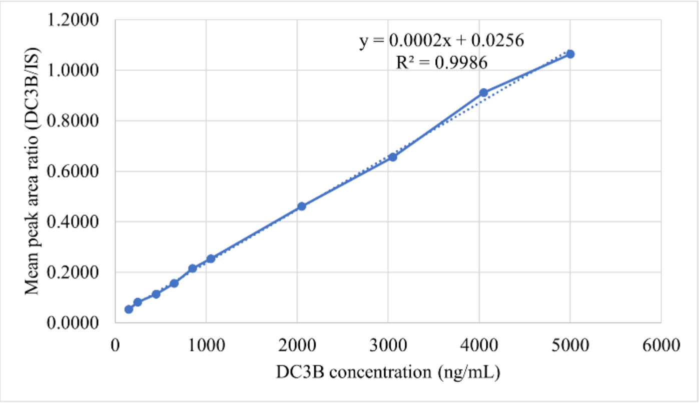

y = 0.0002x + 0.0256
R² = 0.9986
Mean peak area ratio (DC3B/IS)
DC3B concentration (ng/mL)


**I**

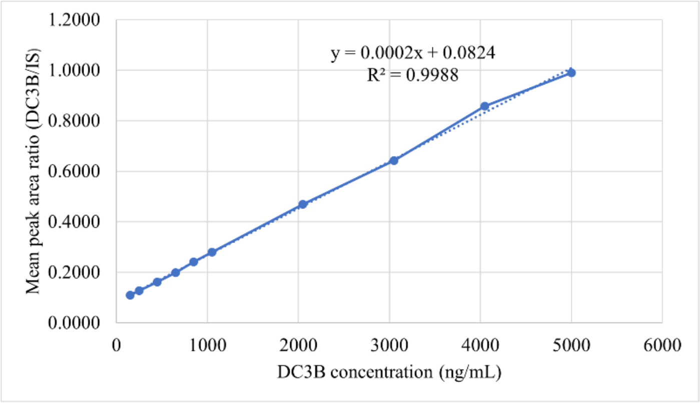

y = 0.0002x + 0.0824
R² = 0.9988
Mean peak area ratio (DC3B/IS)
DC3B concentration (ng/mL)


**J**

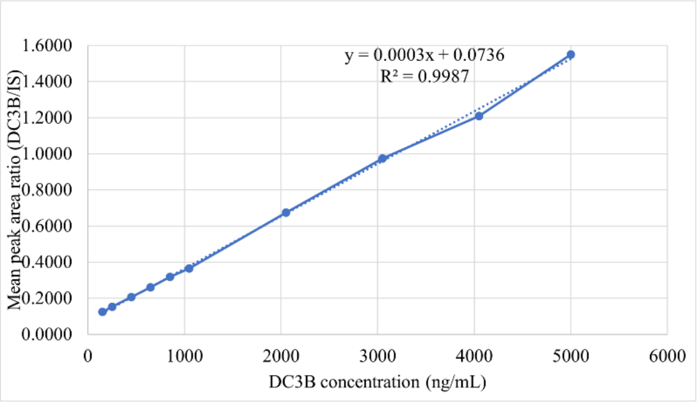

y = 0.0003x + 0.0736
R² = 0.9987
Mean peak area ratio (DC3B/IS)
DC3B concentration (ng/mL)
