## Supporting Information - Tables for "*In vivo* pharmacokinetics and tissue distribution profile of a Wnt/β-catenin pathway-targeting anticancer cassane diterpene isolated from *Caesalpinia pulcherrima*"

**Table S1. Back-calculated concentrations of 6βCHV for plasma and tissue calibration curve samples (n = 3)**

| Tissue matrix | Calibration concentration (ng/mL) | Back calculated concentration (ng/mL) | | RSD (%) | RE (%) |
|---|---|---|---|---|---|
| | | Mean | SD | | |
| **Plasma** | 150 | 138.68 | 3.92 | -7.55 | 2.83 |
| | 250 | 235.94 | 5.77 | -5.62 | 2.44 |
| | 450 | 462.89 | 8.62 | 2.86 | 1.86 |
| | 650 | 674.52 | 7.20 | 3.77 | 1.07 |
| | 850 | 864.05 | 3.51 | -0.46 | 0.42 |
| | 1050 | 1079.81 | 8.62 | 2.84 | 0.80 |
| | 1550 | 1537.10 | 9.16 | -0.83 | 0.60 |
| | 2050 | 2097.64 | 7.67 | 2.32 | 0.37 |
| | 2550 | 2545.89 | 2.66 | -0.16 | 0.10 |
| **Liver** | 150 | 135.29 | 3.82 | -9.81 | 2.82 |
| | 250 | 219.61 | 2.56 | -12.16 | 1.17 |
| | 450 | 427.15 | 4.81 | -5.08 | 1.13 |
| | 650 | 651.61 | 7.94 | 0.25 | 1.22 |
| | 850 | 921.20 | 11.83 | 8.38 | 1.28 |
| | 1050 | 1146.96 | 8.01 | 9.23 | 0.70 |
| | 1550 | 1618.61 | 5.79 | 4.43 | 0.36 |
| | 2050 | 2131.40 | 3.78 | 3.97 | 0.18 |
| | 2550 | 2545.84 | 4.77 | -0.16 | 0.19 |
| **Small intestines** | 200 | 198.77 | 7.46 | 3.75 | -0.61 |
| | 500 | 462.81 | 58.28 | 12.59 | -7.44 |
| | 1000 | 907.91 | 51.24 | 5.64 | -9.21 |

| | | | | | |
|---|---|---|---|---|---|
| | 2000 | 1945.81 | 60.41 | 3.10 | -2.71 |
| | 5000 | 4230.55 | 564.27 | 13.34 | -15.39 |
| | 10000 | 11294.71 | 574.80 | 5.09 | 12.95 |
| | 15000 | 15826.18 | 752.47 | 4.75 | 5.51 |
| | 20000 | 22267.58 | 1156.41 | 5.19 | 11.34 |
| | 25000 | 25747.05 | 569.25 | 2.21 | 2.99 |
| | 30000 | 30446.97 | 466.40 | 1.53 | 1.49 |
| **Stomach** | 200 | 215.02 | 11.26 | 5.24 | 7.51 |
| | 500 | 444.88 | 15.41 | 3.46 | -11.02 |
| | 1000 | 868.70 | 60.89 | 7.01 | -13.13 |
| | 2000 | 1883.11 | 60.95 | 3.24 | -5.84 |
| | 5000 | 4318.68 | 171.95 | 3.98 | -13.63 |
| | 10000 | 11202.83 | 593.89 | 5.30 | 12.03 |
| | 15000 | 14672.70 | 579.74 | 3.95 | -2.18 |
| | 20000 | 21381.90 | 4.47 | 0.02 | 6.91 |
| | 25000 | 22609.73 | 582.63 | 2.58 | -9.56 |
| | 30000 | 28843.79 | 705.08 | 2.44 | -3.85 |
| **Testes** | 150 | 123.41 | 6.87 | 5.56 | -17.73 |
| | 250 | 222.24 | 15.79 | 7.10 | -11.11 |
| | 450 | 473.47 | 47.62 | 10.06 | 5.22 |
| | 650 | 673.69 | 8.92 | 1.32 | 3.65 |
| | 850 | 773.81 | 9.63 | 1.24 | -8.96 |
| | 1050 | 1024.86 | 24.02 | 2.34 | -2.39 |
| | 2050 | 2129.19 | 24.06 | 1.13 | 3.86 |
| | 3050 | 3033.91 | 17.99 | 0.59 | -0.53 |
| | 4050 | 4129.53 | 43.20 | 1.05 | 1.96 |
| | 5000 | 4857.00 | 172.63 | 3.55 | -2.86 |

| | | | | | |
|---|---|---|---|---|---|
| **Lungs** | 150 | 125.58 | 6.25 | 4.98 | -16.28 |
| | 250 | 222.48 | 34.71 | 15.60 | -11.01 |
| | 450 | 475.53 | 7.96 | 1.67 | 5.67 |
| | 650 | 675.84 | 1.72 | 0.26 | 3.97 |
| | 850 | 825.21 | 6.09 | 0.74 | -2.92 |
| | 1050 | 1003.89 | 41.92 | 4.18 | -4.39 |
| | 2050 | 2118.34 | 70.10 | 3.31 | 3.33 |
| | 3050 | 3221.87 | 47.34 | 1.47 | 5.64 |
| | 4050 | 4184.62 | 18.00 | 0.43 | 3.32 |
| | 5000 | 4905.69 | 29.46 | 0.60 | -1.89 |
| **Kidneys** | 150 | 124.83 | 11.00 | 8.81 | -16.78 |
| | 250 | 264.92 | 37.05 | 13.99 | 5.97 |
| | 450 | 402.13 | 42.81 | 10.64 | -10.64 |
| | 650 | 649.37 | 34.64 | 5.33 | -0.10 |
| | 850 | 897.42 | 25.79 | 2.87 | 5.58 |
| | 1050 | 1089.96 | 70.98 | 6.51 | 3.81 |
| | 2050 | 2173.33 | 13.79 | 0.63 | 6.02 |
| | 3050 | 3072.83 | 63.25 | 2.06 | 0.75 |
| | 4050 | 4085.22 | 21.82 | 0.53 | 0.87 |
| | 5000 | 5303.96 | 23.18 | 0.44 | 6.08 |
| **Brain** | 150 | 137.12 | 8.91 | 0.07 | -8.59 |
| | 250 | 280.92 | 8.20 | 0.03 | 12.37 |
| | 450 | 439.48 | 12.67 | 0.03 | -2.34 |
| | 650 | 651.83 | 31.83 | 0.05 | 0.28 |
| | 850 | 944.43 | 52.62 | 0.06 | 11.11 |
| | 1050 | 1139.47 | 143.13 | 0.13 | 8.52 |
| | 2050 | 2177.75 | 67.59 | 0.03 | 6.23 |

| | | | | | |
|---|---|---|---|---|---|
| | 3050 | 3149.91 | 130.08 | 0.04 | 3.28 |
| | 4050 | 4432.02 | 31.54 | 0.01 | 9.43 |
| | 5000 | 5194.37 | 260.70 | 0.05 | 3.89 |
| **Heart** | 150 | 126.58 | 11.76 | 9.29 | -15.61 |
| | 250 | 215.40 | 13.74 | 6.38 | -13.84 |
| | 450 | 386.08 | 14.71 | 3.81 | -14.21 |
| | 650 | 586.02 | 8.52 | 1.45 | -9.84 |
| | 850 | 785.78 | 20.08 | 2.56 | -7.55 |
| | 1050 | 999.39 | 19.81 | 1.98 | -4.82 |
| | 2050 | 1940.09 | 36.77 | 1.90 | -5.36 |
| | 3050 | 2793.69 | 22.99 | 0.82 | -8.40 |
| | 4050 | 3925.67 | 50.26 | 1.28 | -3.07 |
| | 5000 | 4544.58 | 13.99 | 0.31 | -9.11 |
| **Spleen** | 150 | 162.81 | 3.49 | 2.14 | 8.54 |
| | 250 | 268.98 | 11.17 | 4.15 | 7.59 |
| | 450 | 441.06 | 0.62 | 0.14 | -1.99 |
| | 650 | 619.69 | 5.00 | 0.81 | -4.66 |
| | 850 | 814.86 | 3.73 | 0.46 | -4.13 |
| | 1050 | 972.77 | 17.55 | 1.80 | -7.35 |
| | 2050 | 2017.38 | 29.26 | 1.45 | -1.59 |
| | 3050 | 2980.38 | 21.22 | 0.71 | -2.28 |
| | 4050 | 3786.04 | 11.78 | 0.31 | -6.52 |
| | 5000 | 4919.14 | 88.28 | 1.79 | -1.62 |

RE: Relative error; RSD: Relative standard deviation

**Table S2. Intraday and interday accuracy and precision for 6βCHV in rat tissue matrices.**

| Sample matrix | QC level | Nominal concentration (ng/mL) | Intraday (n = 30) | | Interday (n = 25) | |
|---|---|---|---|---|---|---|
| | | | Accuracy (RE%) | Precision (RSD%) | Accuracy (RE%) | Precision (RSD%) |
| **Small intestines** | LLOQ | 200 | 6.21 | 9.68 | 8.45 | 16.96 |
| | LQC | 1000 | -5.03 | 5.75 | -11.25 | 1.07 |
| | MQC | 10000 | 2.20 | 1.10 | 10.81 | 1.19 |
| | HQC | 20000 | 6.68 | 3.23 | 10.48 | 0.49 |
| | ULOQ | 30000 | -8.60 | 2.67 | -15.63 | 0.42 |
| **Stomach** | LLOQ | 200 | 2.01 | 19.05 | 6.52 | 7.97 |
| | LQC | 1000 | -6.71 | 3.22 | -7.41 | 1.08 |
| | MQC | 10000 | 5.08 | 3.46 | 5.79 | 1.55 |
| | HQC | 20000 | 3.21 | 2.19 | 4.60 | 2.86 |
| | ULOQ | 30000 | -0.14 | 1.87 | -1.08 | 1.73 |
| **Testes** | LLOQ | 150 | -8.70 | 4.06 | -6.09 | 0.73 |
| | LQC | 450 | 0.72 | 9.63 | 1.23 | 2.73 |
| | MQC | 1050 | -2.47 | 1.72 | -2.87 | 0.55 |
| | HQC | 3050 | -0.50 | 0.44 | -1.03 | 0.24 |
| | ULOQ | 5000 | -2.76 | 2.81 | -2.63 | 3.23 |
| **Lungs** | LLOQ | 150 | -1.10 | 11.95 | -5.85 | 3.81 |
| | LQC | 450 | 4.36 | 2.11 | 4.96 | 1.85 |
| | MQC | 1050 | -3.37 | 3.26 | -3.46 | 2.10 |
| | HQC | 3050 | 4.02 | 3.19 | -3.20 | 0.03 |
| | ULOQ | 5000 | -0.02 | 1.09 | -3.20 | 1.24 |
| **Kidneys** | LLOQ | 150 | -2.75 | 7.65 | -4.47 | 0.14 |
| | LQC | 450 | -2.53 | 2.15 | -9.09 | 1.67 |
| | MQC | 1050 | 3.03 | 4.75 | 3.99 | 3.15 |

| | | | | | | |
|---|---|---|---|---|---|---|
| | HQC | 3050 | 1.13 | 1.59 | 1.58 | 1.09 |
| | ULOQ | 5000 | 2.33 | 1.72 | 3.02 | 2.49 |
| **Brain** | LLOQ | 150 | -3.30 | 9.91 | 7.44 | 8.90 |
| | LQC | 450 | -3.43 | 3.28 | 3.79 | 4.19 |
| | MQC | 1050 | 2.76 | 1.99 | 6.67 | 4.23 |
| | HQC | 3050 | 2.77 | 3.01 | 2.39 | 2.49 |
| | ULOQ | 5000 | 2.25 | 4.22 | 2.65 | 1.79 |
| **Heart** | LLOQ | 150 | -6.18 | 3.26 | -17.29 | 2.81 |
| | LQC | 450 | -5.48 | 6.17 | -4.18 | 11.25 |
| | MQC | 1050 | -4.86 | 1.60 | 343 | 7.03 |
| | HQC | 3050 | 3.43 | 2.40 | 2.36 | 1.71 |
| | ULOQ | 5000 | -5.55 | 5.17 | -2.22 | 6.86 |
| **Spleen** | LLOQ | 150 | 10.45 | 3.74 | -11.26 | 19.02 |
| | LQC | 450 | -3.16 | 2.43 | 1.86 | 4.85 |
| | MQC | 1050 | -6.38 | 2.06 | 1.21 | 7.05 |
| | HQC | 3050 | -2.28 | 0.71 | 1.59 | 1.52 |
| | ULOQ | 5000 | -1.92 | 1.35 | -7.47 | 14.39 |

QC: Quality control; LLOQ: Lower limit of quantification; LQC: Low quality control; MQC: Medium quality control; HQC: High quality control; ULOQ: Upper limit of quantification; RE: Relative error; RSD: Relative standard deviation

**Table S3. Extraction recovery, extraction efficiency, and matrix effect of 6βCHV in rat liver (n = 6)**

| **Sample matrix** | **QC level/ concentration (ng/mL)** | **Extraction recovery** (%) | | **Extraction efficiency** (%) | | **Matrix effect** (%) | |
|---|---|---|---|---|---|---|---|
| | | **Mean ± SD** | **RSD (%)** | **Mean ± SD** | **RSD (%)** | **Mean ± SD** | **RSD (%)** |
| Small intestines | LQC/1000 | 81.36 ± 3.86 | 4.75 | 75.35 ± 4.58 | 6.07 | 92.66 ± 4.50 | 4.85 |
| | MQC/10000 | 98.22 ± 1.61 | 1.64 | 88.32 ± 1.01 | 1.14 | 89.94 ± 1.85 | 2.06 |
| | HQC/ 20000 | 92.24 ± 3.63 | 3.81 | 93.03 ± 2.51 | 2.70 | 97.73 ± 2.20 | 2.25 |
| Stomach | LQC/1000 | 85.73 ± 5.14 | 6.00 | 75.37 ± 6.09 | 8.07 | 88.01 ± 6.57 | 7.47 |
| | MQC/10000 | 96.79 ± 3.40 | 3.51 | 93.55 ± 2.57 | 2.75 | 96.69 ± 1.61 | 1.66 |
| | HQC/ 20000 | 94.58 ± 3.53 | 3.73 | 89.64 ± 3.18 | 3.55 | 94.80 ± 2.34 | 2.47 |
| Testes | LQC/450 | 85.05 ± 5.15 | 6.06 | 76.05 ± 2.15 | 2.83 | 89.62 ± 4.45 | 4.97 |
| | MQC/1050 | 89.91 ± 3.95 | 4.40 | 80.12 ± 4.89 | 6.10 | 89.22 ± 6.28 | 7.04 |
| | HQC/ 3050 | 96.07 ± 4.80 | 5.00 | 87.80 ± 8.28 | 9.43 | 91.34 ± 5.99 | 6.56 |
| Lungs | LQC/450 | 90.72 ± 3.21 | 3.54 | 80.92 ± 5.19 | 6.42 | 89.20 ± 4.88 | 5.47 |
| | MQC/1050 | 89.18 ± 5.36 | 6.01 | 82.01 ± 6.98 | 8.51 | 91.94 ± 5.04 | 5.48 |
| | HQC/ 3050 | 92.36 ± 5.82 | 6.30 | 87.03 ± 3.32 | 3.82 | 94.41 ± 4.35 | 4.61 |
| Kidneys | LQC/450 | 95.25 ± 6.43 | 6.75 | 83.64 ± 7.99 | 9.56 | 87.73 ± 4.55 | 5.19 |
| | MQC/1050 | 85.60 ± 6.39 | 7.46 | 79.30 ± 6.46 | 8.15 | 92.64 ± 2.96 | 3.19 |
| | HQC/ 3050 | 95.38 ± 3.96 | 4.15 | 88.74 ± 3.94 | 4.44 | 93.05 ± 2.38 | 2.56 |
| Brain | LQC/450 | 8862 ± 4.09 | 4.61 | 76.09 ± 2.41 | 3.17 | 86.00 ± 4.70 | 5.46 |
| | MQC/1050 | 91.05 ± 5.30 | 5.82 | 84.98 ± 4.73 | 5.56 | 93.36 ± 1.93 | 2.07 |
| | HQC/ 3050 | 97.07 ± 2.01 | 2.07 | 94.80 ± 2.28 | 2.41 | 97.66 ± 1.76 | 1.80 |
| Heart | LQC/450 | 96.68 ± 4.18 | 4.18 | 82.45 ± 6.46 | 7.83 | 85.27 ± 5.58 | 6.54 |
| | MQC/1050 | 91.35 ± 3.70 | 4.05 | 84.07 ± 2.67 | 3.17 | 92.07 ± 2.03 | 2.21 |
| | HQC/ 3050 | 95.81 ± 1.05 | 1.10 | 92.94 ± 1.09 | 1.17 | 97.00 ± 1.46 | 1.50 |
| Spleen | LQC/450 | 86.38 ± 6.83 | 7.91 | 78.18 ± 8.36 | 10.70 | 90.41 ± 4.83 | 5.34 |
| | MQC/1050 | 89.27 ± 7.07 | 7.92 | 81.79 ± 4.76 | 5.82 | 91.79 ± 3.76 | 4.10 |
| | HQC/ 3050 | 95.55 ± 3.00 | 3.14 | 92.77 ± 1.66 | 1.79 | 97.13 ± 1.49 | 1.53 |

QC: Quality control; LQC: Low quality control; MQC: Medium quality control; HQC: High quality control; RSD: Relative standard deviation

Table S4.

| Sample matrix | QC level (ng/mL) | Short-term[a] | | Long-term[b] | | Freeze-thaw[c] | | Autosampler[d] | | Post-preparation storage[e] | |
|---|---|---|---|---|---|---|---|---|---|---|---|
| | | RE% | RSD% | RE% | RSD% | RE% | RSD% | RE% | RSD% | RE% | RSD% |
| Small intestines | LQC (1000) | -10.38 | 7.24 | -10.44 | 3.31 | -11.61 | 1.32 | -8.96 | 2.66 | -4.01 | 3.83 |
| | MQC (10000) | -3.20 | 1.01 | -5.59 | 7.27 | -3.06 | 0.29 | -8.04 | 0.67 | -10.59 | 1.05 |
| | HQC (20000) | -3.56 | 0.59 | -6.04 | 0.94 | -9.06 | 0.35 | -10.62 | 0.23 | -6.62 | 0.36 |
| Stomach | LQC (1000) | -7.09 | 2.55 | -11.92 | 2.63 | -1.89 | 0.51 | -10.46 | 9.56 | -4.77 | 2.61 |
| | MQC (10000) | -3.12 | 0.67 | -2.33 | 0.76 | -3.63 | 0.39 | -7.98 | 0.87 | -12.82 | 0.53 |
| | HQC (20000) | -2.96 | 0.47 | -1.39 | 0.48 | -6.95 | 0.35 | -9.05 | 0.39 | -8.06 | 0.26 |
| Testes | LQC (450) | -4.13 | 3.41 | -6.48 | 4.53 | -11.30 | 5.61 | -5.53 | 5.35 | -5.82 | 1.54 |
| | MQC (1050) | -5.22 | 2.82 | -7.13 | 4.21 | -8.09 | 1.79 | 7.94 | 3.44 | -8.43 | 2.22 |
| | HQC (3050) | -4.84 | 2.67 | -2.51 | 1.29 | -5.74 | 0.53 | -6.51 | 1.19 | -6.18 | 1.00 |
| Lungs | LQC (450) | -7.51 | 3.36 | -5.58 | 7.78 | -12.10 | 5.87 | -11.97 | 1.48 | -11.11 | 2.65 |
| | MQC (1050) | -6.96 | 1.24 | 1.43 | -4.89 | -6.49 | 3.89 | -6.64 | 2.32 | -6.27 | 1.85 |
| | HQC (3050) | -0.87 | 0.52 | 1.31 | -6.59 | -8.38 | 3.50 | -3.73 | 1.25 | -2.83 | 1.48 |
| Kidneys | LQC (450) | -6.54 | 1.47 | -6.29 | 2.11 | -10.00 | 3.64 | -9.41 | 3.51 | -8.09 | 2.41 |
| | MQC (1050) | -3.41 | 1.20 | -2.61 | 1.72 | -2.98 | 3.54 | -4.39 | 1.75 | -3.38 | 1.25 |
| | HQC (3050) | -1.33 | 0.33 | -1.76 | 1.01 | -6.22 | 0.79 | -12.89 | 1.14 | -13.15 | 1.20 |
| Brain | LQC (450) | -5.85 | 1.82 | -12.11 | 2.61 | -9.54 | 4.20 | -12.49 | 2.84 | -12.08 | 317 |
| | MQC (1050) | -1.74 | 0.42 | -3.66 | 2.18 | -3.74 | 2.15 | -5.15 | 1.52 | -6.10 | 2.25 |
| | HQC (3050) | -0.64 | 0.40 | -5.89 | 1.14 | -6.14 | 2.40 | -3.05 | 0.88 | -6.26 | 1.57 |
| Heart | LQC (450) | -9.64 | 6.20 | -12.44 | 1.94 | -11.65 | 3.97 | -11.67 | 1.94 | -9.52 | 2.23 |
| | MQC (1050) | -2.59 | 0.79 | -6.60 | 1.89 | -4.90 | 2.25 | -4.84 | 1.36 | -4.87 | 1.36 |
| | HQC (3050) | -1.21 | 0.24 | -3.30 | 1.20 | -9.46 | 1.17 | -7.03 | 0.80 | -6.57 | 1.17 |
| Spleen | LQC (450) | -8.45 | 2.32 | -12.27 | 2.48 | -11.51 | 6.84 | -13.52 | 2.08 | -15.34 | 2.47 |
| | MQC (1050) | -3.43 | 1.29 | -6.27 | 0.74 | -4.22 | 3.24 | -5.35 | 1.51 | -6.22 | 2.18 |
| | HQC (3050) | -1.33 | 0.41 | -6.67 | 1.09 | -1.25 | 1.09 | -2.70 | 1.16 | -2.35 | 1.45 |

[a] Plasma and liver for 6h at room temperature. [b] Plasma and liver for 30 days at -20 ºC and -80 ºC, respectively. [c] Three freeze-thaw cycles from -80 ºC to room temperature for liver and -20 ºC to room temperature for plasma. [d] Plasma and liver samples were stored in the autosampler for 24

h before injection. [e] Injection-ready plasma and liver QCs at 4 ºC for 48 h. LQC: Low quality control, MQC: Medium quality control, HQC: High quality control

**Table S5. Dilution integrity experiment of DC3B in rat plasma and tissue matrices. Dilution quality control samples were diluted 2-fold (n = 5, mean ± SD)**

| Matrix/ ULOQ (ng/mL) | Pre-diluted concentration (ng/mL) | Diluted concentration (ng/mL) | Measured concentration (ng/mL) | RE (%) | RSD (%) |
|---|---|---|---|---|---|
| **Plasma/ 2550** | 5100 | 2550 | 2299.81 ± 32.39 | -9.81 | 1.41 |
| | 3000 | 1500 | 1325.49 ± 56.47 | -11.63 | 4.26 |
| **Liver/ 2550** | 5100 | 2550 | 2369.15 ± 26.15 | -7.09 | 1.10 |
| | 3000 | 1500 | 1314.41 ± 30.45 | -12.37 | 2.32 |
| **Stomach/ 30000** | 60000 | 30000 | 29463.38 ± 321.95 | -1.79 | 1.09 |
| | 40000 | 20000 | 19412.78 ± 274.94 | -2.94 | 1.42 |
| **Small intestines/ 30000** | 60000 | 30000 | 29367.84 ± 298.16 | -2.11 | 1.02 |
| | 40000 | 20000 | 19338.11 ± 312.79 | -3.31 | 1.62 |
| **Testes/ 5000** | 10000 | 5000 | 4678.91 ± 54.66 | -6.42 | 1.17 |
| | 7000 | 3500 | 3249.11 ± 42.13 | -7.17 | 1.30 |
| **Lungs/ 5000** | 10000 | 5000 | 4865.10 ± 13.27 | -2.70 | 0.27 |
| | 7000 | 3500 | 3279.64 ± 10.07 | -6.30 | 0.31 |
| **Kidneys/ 5000** | 10000 | 5000 | 4655.09 ± 52.61 | -6.90 | 1.13 |
| | 7000 | 3500 | 3161.47 ± 43.92 | -9.67 | 1.39 |
| **Brain/ 5000** | 10000 | 5000 | 4412.34 ± 45.65 | -11.75 | 1.03 |
| | 7000 | 3500 | 3109.37 ± 43.92 | -11.16 | 2.17 |
| **Heart/ 5000** | 10000 | 5000 | 4722.94 ± 32.14 | -5.54 | 0.68 |
| | 7000 | 3500 | 3101.19 ± 17.82 | -11.39 | 0.57 |
| **Spleen/ 5000** | 10000 | 5000 | 4313.71 ± 45.46 | -13.73 | 1.05 |
| | 7000 | 3500 | 3287.73 ± 41.94 | -6.06 | 1.28 |

ULOQ: Upper limit of quantification; RE: Relative error; RSD: Relative standard deviation
